# What’s so special about Giardia ventral flagella? Interspecies cross-reacting monoclonal antibody against *Pneumocystis jiroveci* reacts with cilia and sparks questions on host-pathogen interactions

**DOI:** 10.1101/2020.05.09.085829

**Authors:** Ewert Linder

**Affiliations:** Swedish Institute for Infectious Disease Control, Solna, Sweden; Department of Microbiology, Tumor and Cell Biology (MTC), Karolinska Institutet, Solna, Sweden

## Abstract

A mouse monoclonal antibody (Moab 4B8) cross-reacting with cilia/flagella was obtained by immunization with Pneumocystis-infected human lung tissue. A key observation was that Moab 4B8 reacted with the ventral flagella of *Giardia intestinalis*, but not with the three other flagellar pairs of this protozoan. To further identify the 4B8 target, its distribution was studied by immunofluorescence staining of cells and tissues of various origin.

The target epitope recognized by Moab 4B8 was found to be associated with structures rich in microtubules; e.g. the mitotic spindle of cultured cells, ciliated airway epithelia, Sertoli cells of the testis and ependymal cells lining brain ventricles. The conserved nature of the 4B8 target was further shown by its presence in cilia of metazoan Schistosome larva and the green alga *Chlamydomonas reinhardtii*. Absence of the 4B8 target from Trypanosomes and Leishmania flagella suggested that it is involved in some function not primarily related to motility. Its presence in only the ventral flagella of Giardia therefore provides a unique opportunity to elucidate the relationship between ciliary structure and function in the same organism.

The observed locations of the 4B8 target in tissues and cells of various origin, suggest a similarity to annexins - and specifically to α-19-giardin. This raises the possibility that it is involved in intra-flagellar transport and provides a basis for further studies aiming at its identification.

**Author Summary:** Pneumocystis is a ubiquitous fungal organism apparently colonizing the lung at an early age to cause pneumonia only in individuals with an impaired immune system. In the alveolar spaces of such individuals, extensive and frequently fatal proliferation of the pathogen occurs. Pneumocystis has no known reservoir in nature and apparently is transmitted directly from infected individuals via an airborne route. Adaptation of this Ascomycotic fungus to a parasitic lifestyle during its evolution apparently resulted in dependence upon host nutrients, but little is known about this presumed adaptation process. In this report, a previously unrecognized constituent of human Pneumocystis is detected using a monoclonal anti-*Pneumocystis jiroveci* antibody (Moab 4B8) which was obtained as a by-product in the search for reagents useful in diagnostics. The Moab 4B8 was shown to react with Pneumocystis but also with cytoskeletal microtubules, e.g. in ciliated epithelia, but not ubiquitously a constituent of the conserved cilia/flagella axonemal structure. A striking example of the discriminating capacity of antibody 4B8 was seen in immunofluorescent staining of the protozoan *Giardia intestinalis*, where only one out of four flagellar pairs expresses the target epitope. This observation of flagellar heterogenicity provoked the question raised in the title of this report. It also provides the basis for the discussion, which arrives at suggestive evidence for the involvement of the described evolutionarily conserved target in host-pathogen interactions related to membrane transport.

## Background

At the onset of the aids epidemic we attempted to develop improved diagnostics of Pneumocystis pneumonia by generating monoclonal antibodies to Pneumocystis (1)(2). Massive clusters of intra-alveolar Pneumocystis organisms causing often fatal pneumonia in infected lungs (3) was a re-emerging problem associated with immunosuppression as pneumocystis pneumonia was known to be fatal in malnourished children in post-war Europe.(4)(5) Cysts could be identified by the pathologist using silver stains such as the Grocott-Gomori silver precipitation stain commonly used to visualize fungi, but higher sensitivity could be achieved using monoclonal antibodies as immuno-cytological markers. (1)(2) With the recognition that Pneumocystis is a fungal organism and the identification of the host-specific human pathogen *Pneumocystis jiroveci*. (6), the cysts are regarded as asci of Ascomycete fungi and the ‥trophozoites‥ equivalent to vegetative yeast.

Monoclonal antibodies, such as 3F6, which were developed to improve the sensitivity of diagnostics were shown to detect not only the cyst form, but also the trophozoites (1), which are the main constitutes of intra-alveolar clusters of organisms in pneumocystis pneumonia (7) (8). However, some of the anti-Pneumocystis monoclonal antibodies, like 4B8, obtained by immunization with a urea extract of Pneumocystis infected human lung tissue also reacted with controls such as cultured mammalian cells and tissues and therefore regarded as useless. Especially the recently recognized fundamental importance of cilia, motivated an attempt to further explore the significance of the observed cross reaction of antibody 4B8 with the ventral flagella of *Giardia intestinalis* (9).

## Materials and Methods

### Pneumocystis materials and generation of anti-Pneumocystis antibodies

Crude extracts of Pneumocystis infected lung tissue, obtained at autopsy from a fatal case of Pneumocystis pneumonia (PCP) was prepared by homogenization in 8M urea, followed by centrifugation at 1000 g. The supernatant, was used for immunization of CBA mice for the production of monoclonal antibodies (Moab) as described earlier (1).

Briefly, hybridoma culture supernatants were screened for antibody activity by two assays: An enzyme-immunoassay (ELISA) in which microtiter plates were coated with the 8 M urea lung extract used for immunization (protein concentration of 10 ng/ml.) Bound antibodies were detected with horseradish-peroxidase-(HRP) conjugated sheep anti mouse immunoglobulin conjugate followed by substrate/chromogen solution which was recorded with a Titer-Tec (Flow Laboratories) fotometer.

ELISA-reactive supernatants were then tested for reactivity by an indirect immunofluorescence (IFL) assay using paraffin sections of Pneumocystis infected lung tissue as antigen (see below).

Electron microscopy was performed on ultrathin sections of Pneumocystis-infected lung material using standard fixation and epon-embedding techniques essentially as described before (10) (11)

ELISA and IFL-reactive hybridoma supernatants were tested by Western blotting after separation of the urea-soluble tissue material by polyacrylamide gel electrophoresis (PAGE) (12) and electrophoretic transfer of material on to nitrocellulose paper (13). The sheep anti-mouse HRP conjugate was used in concentration of 10 mg/ml.

Molecular weight standards were: carbonic anhydrase 30, ovalbumin 43, albumin 67 kDa, phosphorylase b 94 kDa. (Pharmacia, Sweden)

A two-cycled microtubule protein preparation of bovine brain (14) containing mainly alpha and beta tubulins (approximately 80%), kindly provided by Margareta Wallin, University of Gothenburg, was used as reference in Western blotting experiments.

### Cells, tissues and parasites used as antigens. Antibodies used in immunolocalization experiments

Vero cells and Hela cells were cultured under standard conditions in RPMI 1640 medium containing 10% foetal calf serum glutamine, penicillin and streptomycin Flow Laboratories, Herts, England. Cells grown on glass coverslips or suspensions of cells were smeared as monolayers on microscope slides and fixed in cold acetone as described previously (15).For Giardia intestinalis WB isolate in axenic culture we used TYI-S-33 medium supplemented with bile. (16) *Chlamydomonas reinhardi* algae in Sueoka’s medium (17) were provided by Susan Dutcher, Department of Genetics, Washington University School of Medicine, St. Louis, Missouri.

Vinblastine sulphate (Sigma Chemical Co, St Louis, Mo) was added to cell culture medium to disrupt the microtubules cytoskeleton (18)

Different life cycle stages of *Schistosoma mansoni* worms were obtained from parasite life cycle maintained in the laboratory as described previously (19). For immunohistology, human lung tissue, mouse tissues and *Schistosoma mansoni* worms were fixed in Bouin’s fixative (5% acetic acid 9% formaldehyde, 0.9% picric acid) and embedded in paraffin for preparation of tissue sections for immunohistology as described previously (10). Monoclonal antibodies (Moab) 3F6, 4B8 and 2E3 against Pneumocystis were generated as described above (1). Moab 3F6 was also obtained from DAKO/Agilent CODE IR635 (https://www.agilent.com/cs/library/catalogs/public/00230_atlas_of_stains.pdf). Monoclonal anti-tubulin antibodies 1A2, 3G6 and rabbit antibody antiTB ASP were from Thomas Kreis† EMBL, Heidelberg (20).

Deparaffinized tissue sections and fixed cultured cells on microscope slides were incubated with hybridoma culture supernatant, to detect bound antibodies, a sheep anti-mouse immunoglobulin FITC conjugate (National Bacteriology Laboratory (NBL), Stockholm) was used. Fluorescence microscopy was performed using a Zeiss standard microscope. Some microscope pictures from double stained tissue sections obtained on Kodak Tri-X black and white negative film were pseudo-colored using Adobe Photoshop CC 2019 graphics editor. Some of the immunohistology and TEM preparations containing Pneumocystis material was made available for virtual microscopy (21) at the Webmicroscope home page: http://demo.webmicroscope.net/research/parasitology.

## Results

The 4B8 antibody recognizes a Pneumocystis surface antigen cross reacting with a variety of tissues and organisms. Its distribution was similar or identical to that of reference anti-tubulin antibodies e.g in cultured cells. However, co-localization with tubulin was not absolute: The most striking example was the selective localization of the epitope at the ventral but not the anterior, posterior-lateral or caudal flagellar pair of *Giardia intestinalis*.

Thus, the 4B8 target epitope, was not seen in all axonemal structures and co-localization with tubulin was not universal.

### Reaction of antibody 4B8 with Pneumocystis and cross-reactivity by indirect immunofluorescence with protozoa, algae and metazoa

Lung tissue used for preparation of the material for immunization and generation of mouse monoclonal antibodies contained abundant Pneumocystis organisms, as seen in both light and electron micrographs. (Fig.1)

**Fig. 1.**
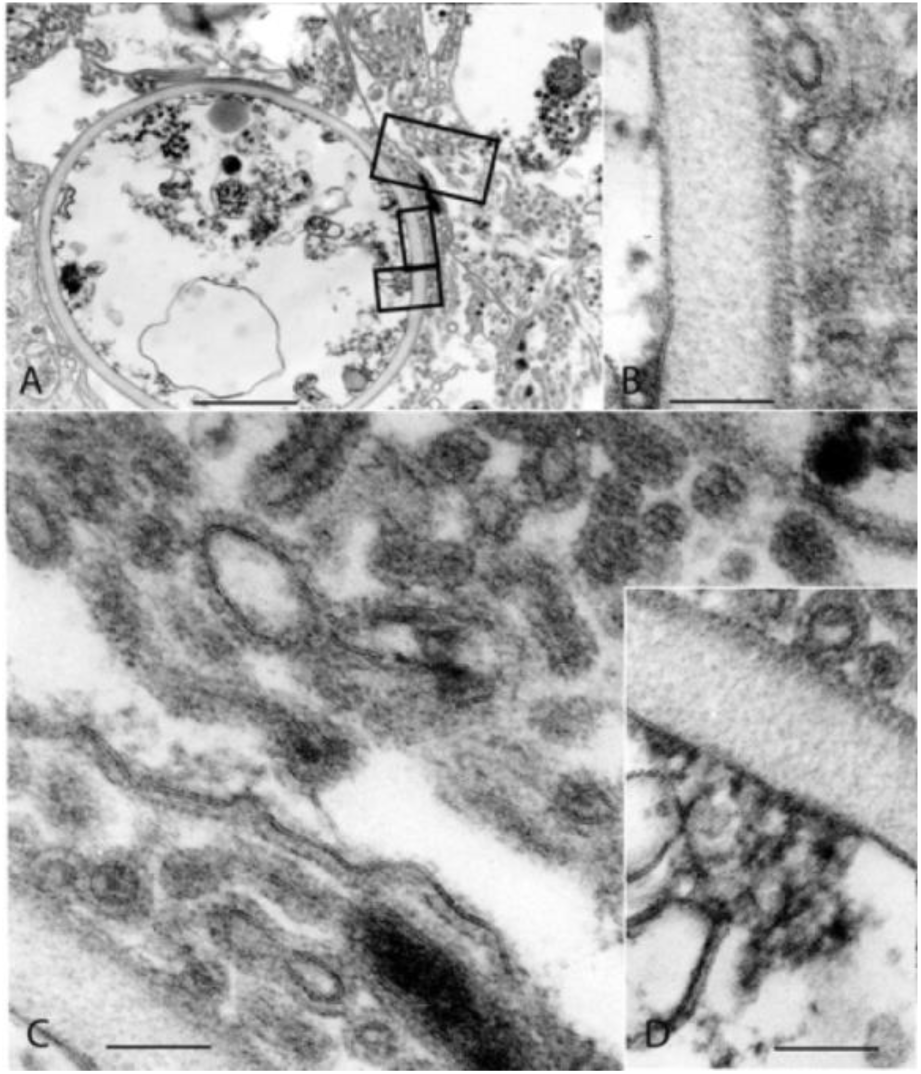
Ultrastructural appearance of *Pneumocystis jiroveci* organisms in lung tissue from fatal case of pneumocystis pneumonia used in this study for the generation of monoclonal antibodies. Cross section of an ascus (cyst) ***(A)***. In inserts ***(B, C, D)***, three morphologically distinct structures are indicated: Intra-cystic trophic forms (tr), tubular extensions (tu) (^‥^filopodia^‥^), which are continuous with the outer electron dense layer of the wall of the ascus. Bar equals 1 micron.

The antibody 4B8 stained *Pneumocystis jiroveci* cysts (asci) in intra-alveolar parasite clusters similarly as the parasite-specific antibody 3F6. Clusters of Pneumocystis organisms can be distinguished in paraffin sections stained by indirect immunofluorescence (Fig 2) The abundance of intra-alveolar Pneumocystis organisms is evident in tissue sections, which can be examined by virtual microscopy and additional ultrastructural observations can be seen at http://demo.webmicroscope.net/research/parasitology. Reference anti-tubulin antibody 1A2 stained intra-cystic material but not the cyst wall. (Fig. 2)

**Fig. 2.**
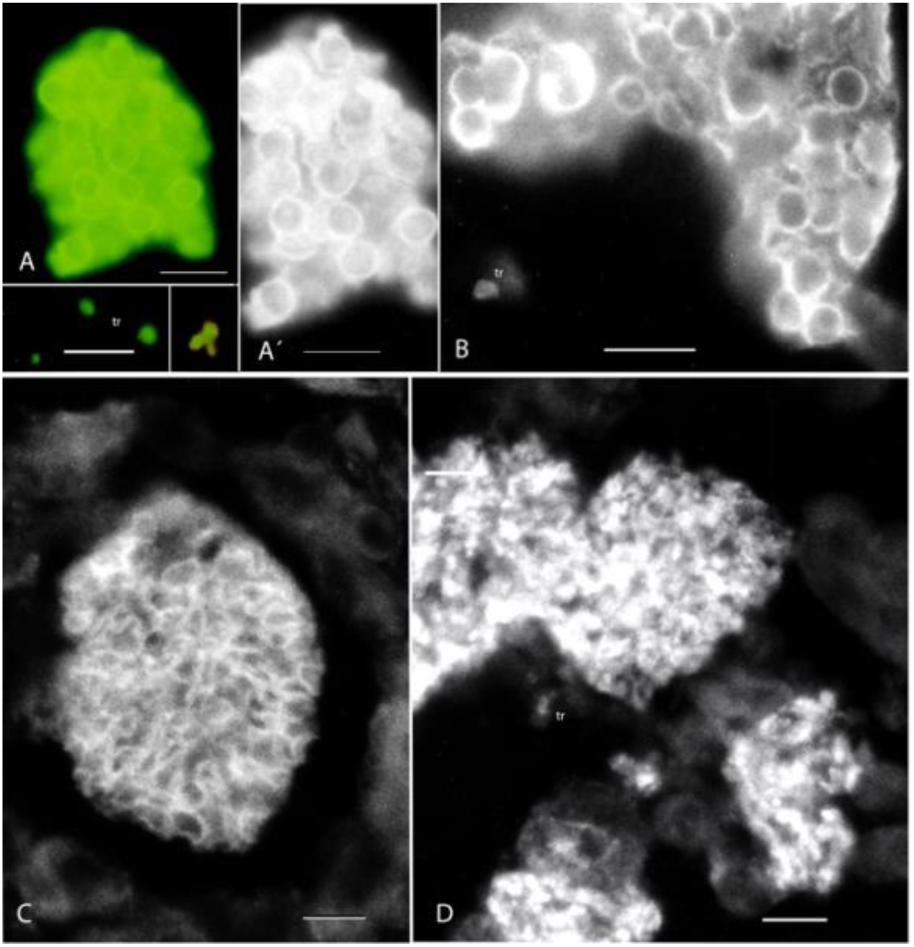
In broncho-alveolar lavage samples from patient with pneumocystis pneumonia, clusters of Pneumocystis organisms are stained similarly by indirect immunofluorescence with monoclonal antibody 4B8 ***(A and A’)*** and parasite-specific antibody3F6 ***(B)***. Note staining of both cyst walls and material occupying the space between cysts. Both antibodies also react with trophozoites (tr). In paraffin sections of lung tissue, cyst walls of intra-alveolar Pneumocystis clusters stain distinctly with 4B8 ***(C)*** whereas anti-tubulin antibody 1A2 gave a lumpy or coarse granular reaction pattern, but no staining of cyst walls ***(D)***. Bar equals 15 microns

By immunoblotting, antibody 4B8 reacted with 3 components, 82, 60 and 45 kDa components present in the Pneumocystis extract whereas reference antibody 3F6 like reacted with only one 82kDa component. In Giardia preparations, a 45-50 kDa component was seen. 4B8 reacted with a component slightly smaller than the tubulin band recognized by Moab 1A2 in the bovine brain tubulin preparation. (Fig. 3)

**Fig. 3.**
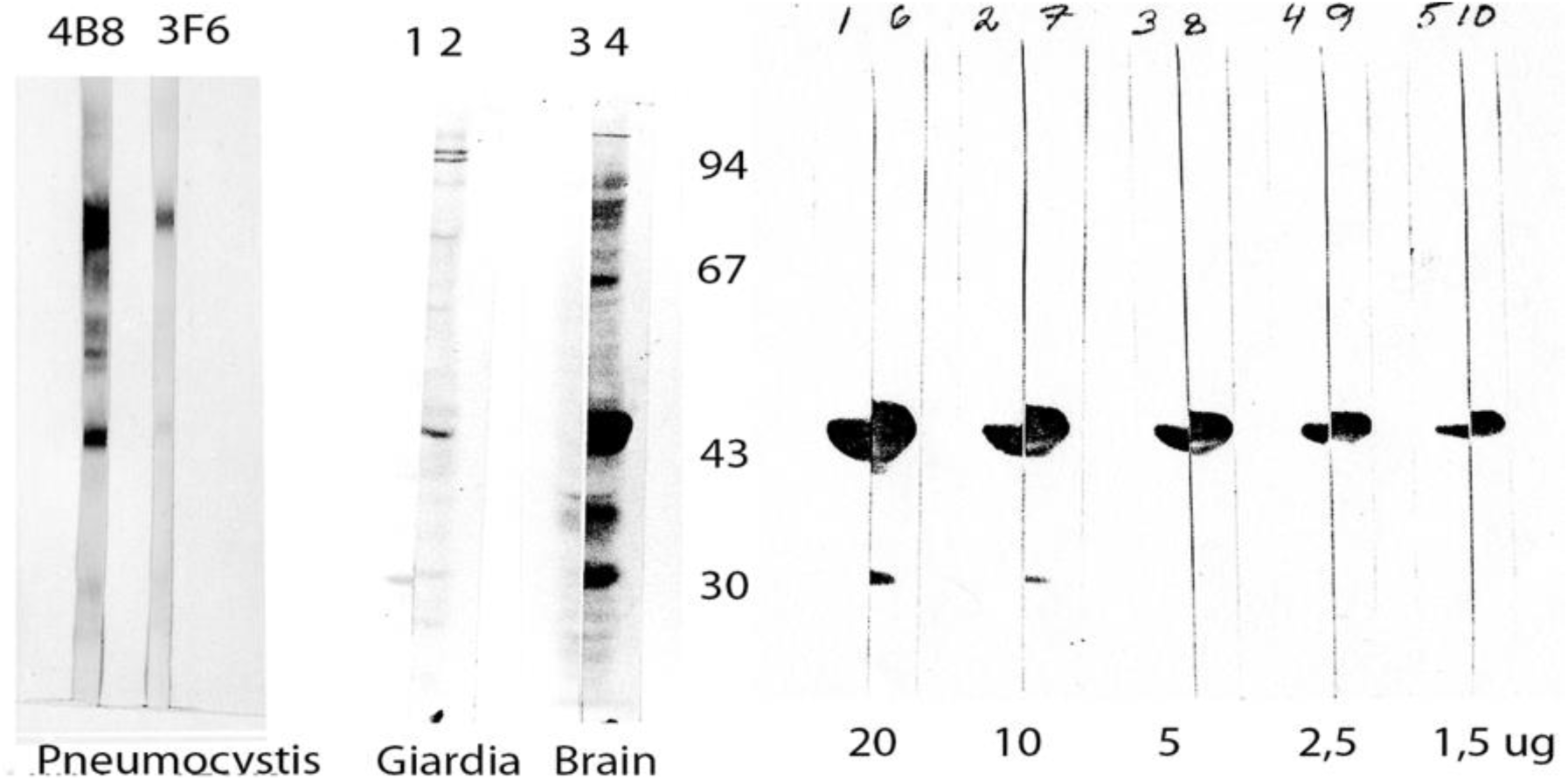
Immunoblotting using monoclonal antibodies 4B8 and control Pneumocystis-specific antibody 4B8 on nitrocellulose paper transfer of *Pneumocystis-*infected lung tissue urea-extract after one-dimensional polyacrylamide gel electrophoresis. Reactivity of 4B8 with major Pneumocystis components of about 80 kDa and45 kDa. 3F6 reacts with compoinent of about 80 kDa. Reactivity of Giardia and rat brain with antibody 4B8. Control strips (lanes 1 and 3) are buffer controls not incubated with monoclonal antibody. Reactions with bovine brain tubulin in amounts from 1.5 to 20 ug. Antibody 4B8 reacts with a component (lanes 1-5) which is slightly smaller than the component reacting with anti-tubulin antibody 1A2 (lanes 6-10). Size markers: Phosphorylase B (94 kDa); human serum albumin (67 kDa); bovine serum albumin (43 kDa) and carbonic anhydrase (30 kDa).

By indirect immunofluorescence staining, antibody 4B8 reacts with some but not all flagella. This was seen as reactivity with some but not all flagellated organisms. The flagella of Chlamydomonas reacted (Fig. 4), whereas *Trypanosoma cruzi* and *Leishmnania tropica* flagella did not (Fig 5).

**Fig. 4.**
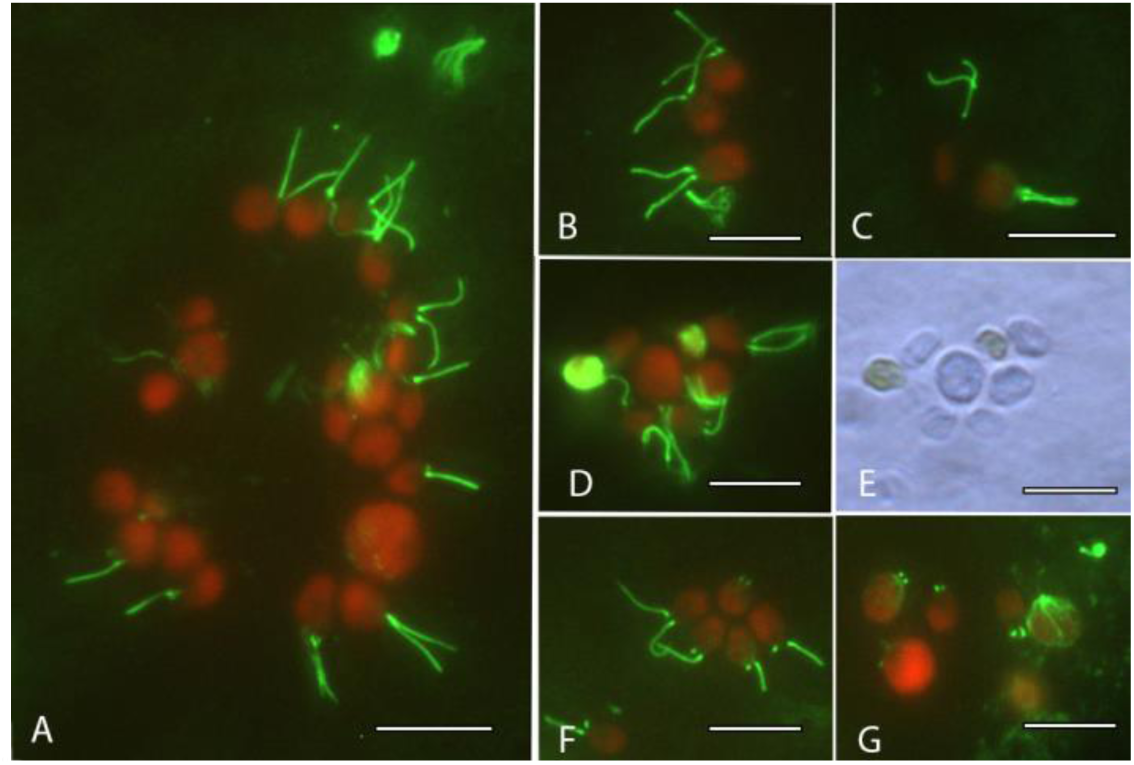
Reactivity of monoclonal antibody 4B8 with *Chlamydomonas reinhardti* flagellar pair and basal bodies. Note staining of the basal bodies also in organisms which have lost one or both flagella during the staining procedure. (A, F and G) Redistribution of the 4B8 target antigen during cell division is seen in G and D/E. Bars equal 10 microns

**Fig. 5.**
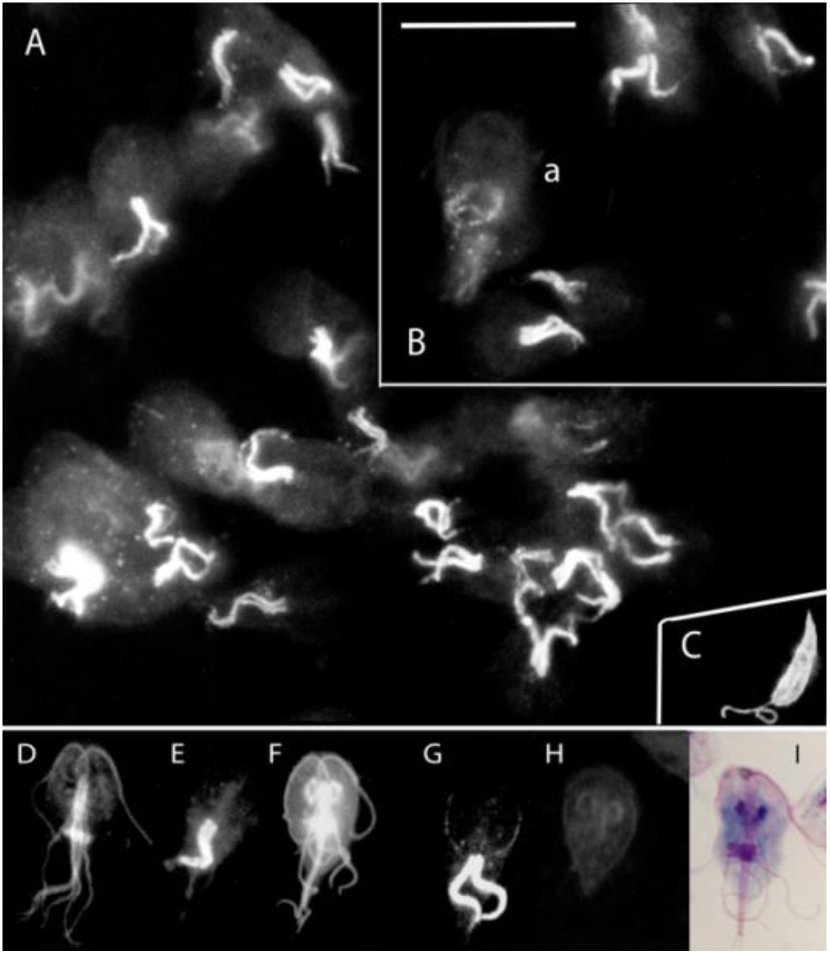
Reactivity of monoclonal antibody 4B8 with *Giardia intestinalis* ***(A, B, E*** and ***G)***.Anti-tubulin antibody 1A2 with. 4B8 reacts only with the ventral flagellar pair. Anti-tubulin reference antibody reacts with all Giardia flagella and in *Leishmnania tropica* epimastigotes both the flagellum and the cell membrane ***(C)*** Monoclonal anti-tubulin antibodies 1A2 ***(D)*** and 3B6 ***(F)*** react with all flagella and the median body. Rabbit antibody antiTB ASP shows weak perinuclear reaction. Bar equals 10 microns

The flagella of these protozoa, however, showed reaction with anti-tubulin antibodies (Fig 5C).

Reactive and non-reactive flagella were present in the same organism, *Giardia intestinalis*. The Giardia ventral flagellar pair reacted, but not the 3 other bilaterally symmetrical pairs, the anterior, posterior-lateral and caudal ones (Fig. 5). In some dividing cells, intracytoplasmic fine granular strands were seen (Fig 5Ba)

Antibody 4B8 appeared not to stain the intracytoplasmic part located of the posteriorly directed ventral flagella. Control staining with anti-tubulin antibodies showed reactivity with all Giardia flagella.

In the metazoan parasitic worm *Schistosoma mansoni*, antibody 4B8 reacted with three distinct cell types; the ciliated surface epithelium of miracidia, flame cells and the nervous system. (Fig. 6).

**Fig. 6.**
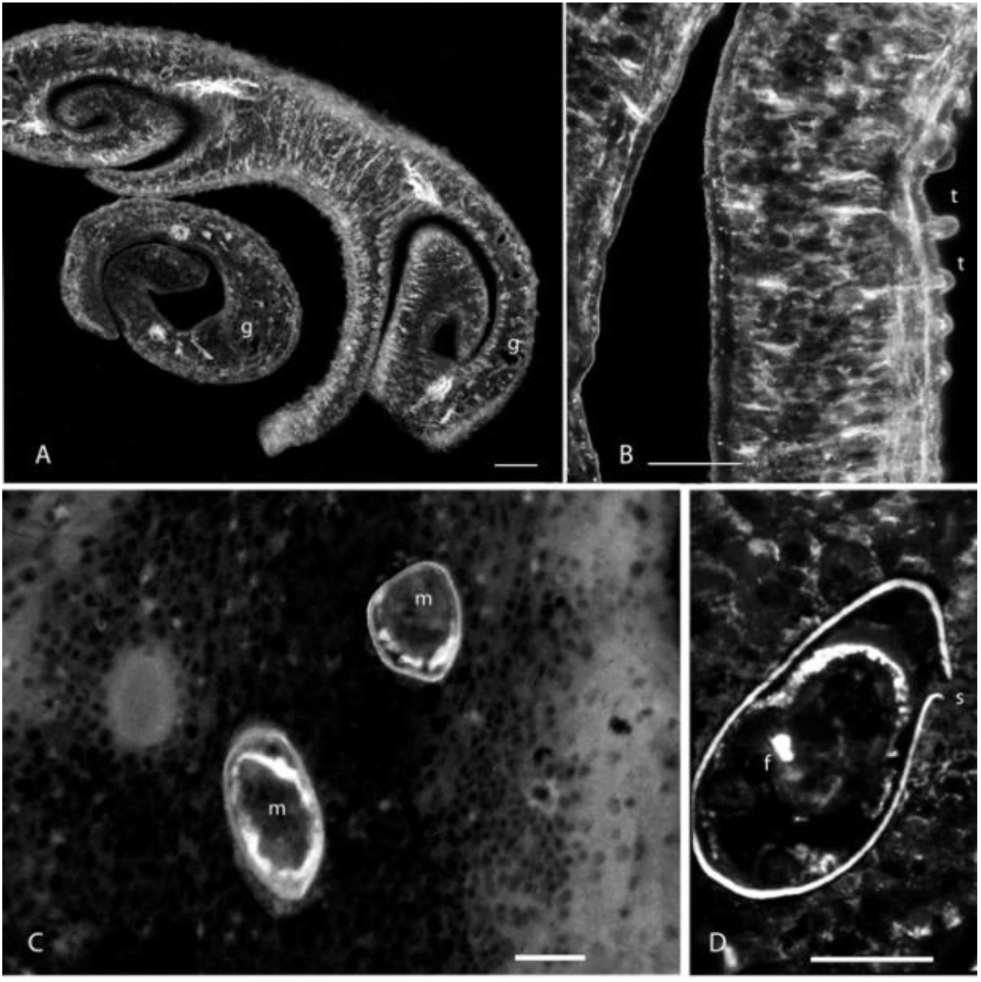
Reactivity of monoclonal antibody 4B8 with *Schistosoma mansoni* adult worm pair. ***(A)***, *Schistosoma mansoni* male worm ***(B)*** shows localization in the longitudinal nerve bundles and in subtegumental nerve fibers. The ciliated surface (m) and flame cell (f) of intraoval miracidia ***(C)*** and ***(D)***. Bars equal 50 microns

In cultured fibroblasts and HeLa cells the 4B8 TA co-localized with tubulin and was present in the mitotic spindle. In cell cultures treated with the microtubulus-disrupting drug Vinblastine, the 4B8 TA was seen in cytoplasmic paracrystals (Fig. 7),

**Fig. 7.**
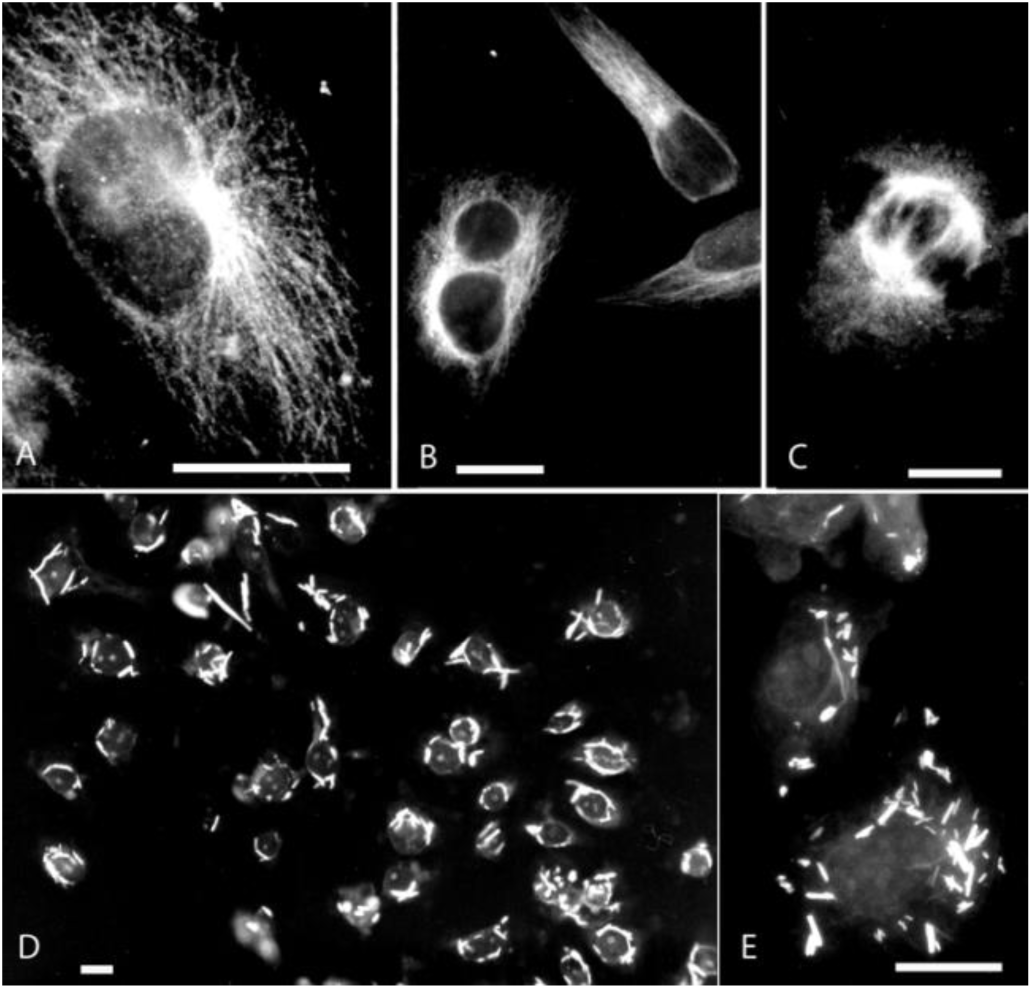
Reactivity of monoclonal antibody 4B8 with cultured Vero cells ***(A)***. Similar staining pattern is seen with anti-tubulin 1A2 ***(B)***. Staining using antibody 4B8 of the mitotic spindle ***(C)*** and staining of paracrystals in cells in after addition of microtubule-disrupting drug Vinblastine to the culture medium ***(D, E)***. Bars equal 5 microns.

### Reaction of antibody 4B8 with tissues

In rat tissues (Fig. 8), Moab 4B8 reacted with ciliated epithelia in several tissues studied, e.g. bronchial epithelium, Fallopian tubes and ependymal cells. The 4B8 TA was seen in both central and peripheral nervous system. In some cells without cilia such as cells in the central nervous System both in the brain cortex (Fig. 8E) and in Purkinje cells of the cerebellum (Fig. 8D) the staining pattern appeared to be granular.

**Fig. 8.**
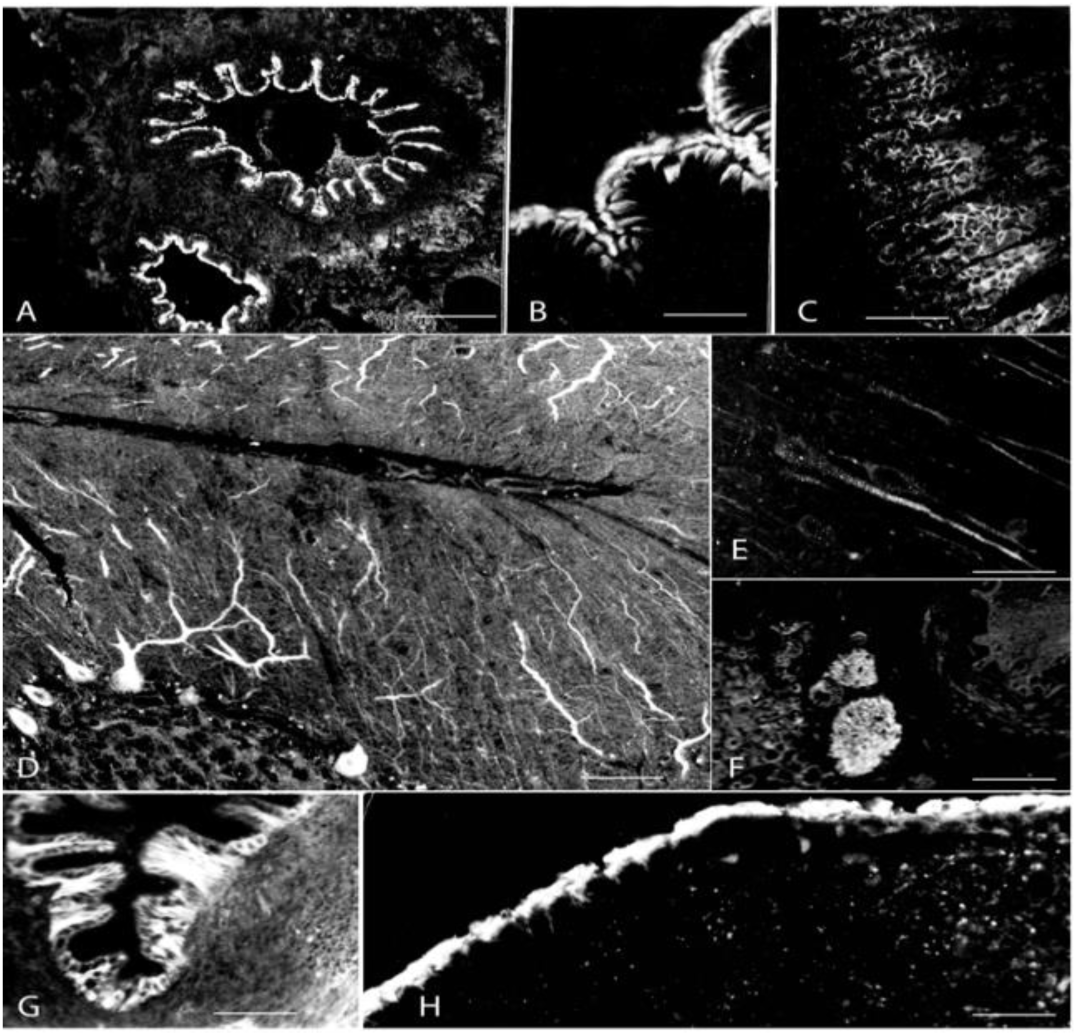
Reactivity of monoclonal antibody 4B8 with mouse tissues. Epithelium of bronchus ***(A)*** and oesophagus ***(B)***. Gastric epithelium ***(C)***. Purkinje cells of the cerebellum ***(D)***. Nerve cells in the cerebral cortex-note the fine granular staining pattern ***(E)***. Peripheral nerves ***(F)***. Fallopian tubes ***(G)***. Ependymal cells ***(H)***. Bars equal 40 microns

The 4B8 target antigen was present in the seminiferous tubes of the testis in Sertoli cells and developing spermatocytes (Fig. 9).

**Fig. 9.**
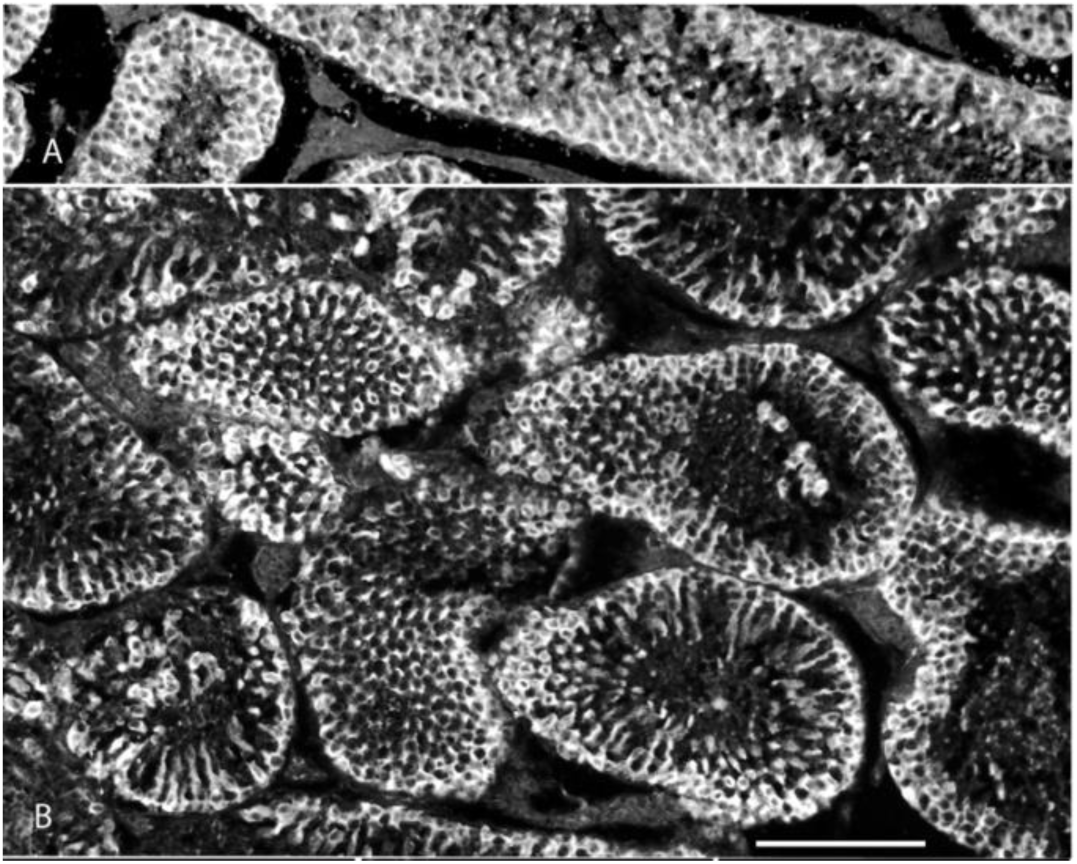
Reactivity of monoclonal antibody 4B8 with rat testis. Seminiferous tubes are seen in longitudinal ***(A)*** and transverse sections ***(B)***. Both developing spermatocytes and Sertoli cells react, whereas interstilial areas containing Leydig cells do not. Bar equals 150 microns.

**Fig. 10.**
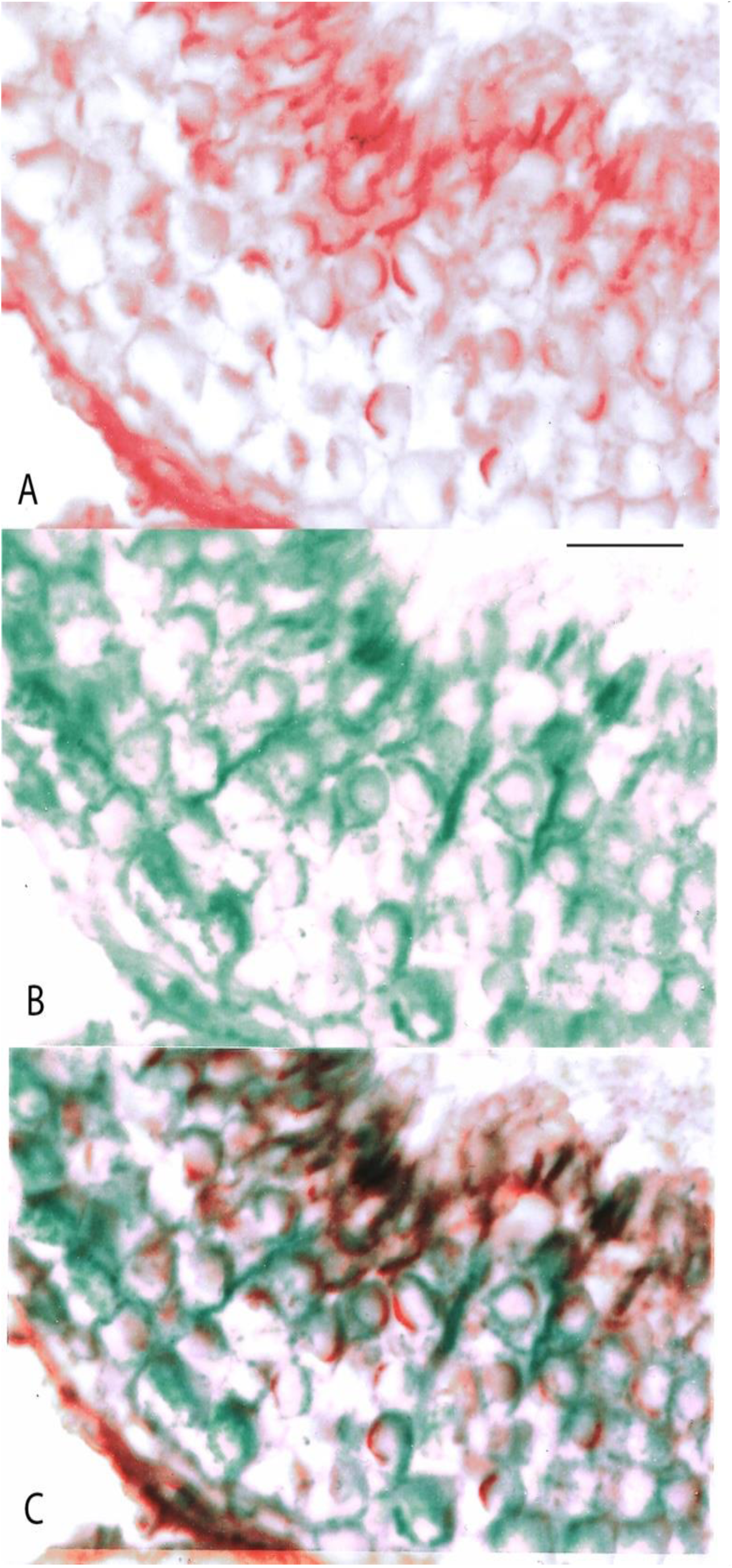
Staining by fluorescence microscopy of the sperm head acrosome with fluorochrome labeled wheat germ agglutinin lectin (WGA) (A) Staining by indirect immunofluorescence of developing spermatocytes and Sertoli cells with mouse monoclonal antibody 4B8 ***(B)***. Partial co-distribution of the two staining patterns is seen in overlay of the WGA (red) and 4B8 (green) staining patterns of the same double-stained tissue section. An overlay of WGA and 4B8 is seen in ***C***. Note that opposite areas of some spermatocyte nuclei are stained red or green, whereas overlapping areas are seen as brown especially in the seminiferous epithelium close to the lumen in the upper part of figures. Figures were obtained from Kodak TriX black and white negative film by pseudo-coloring of original photographs obtained using Adobe PhotoshopR. Bar equals 20 microns.

**Table 1.**
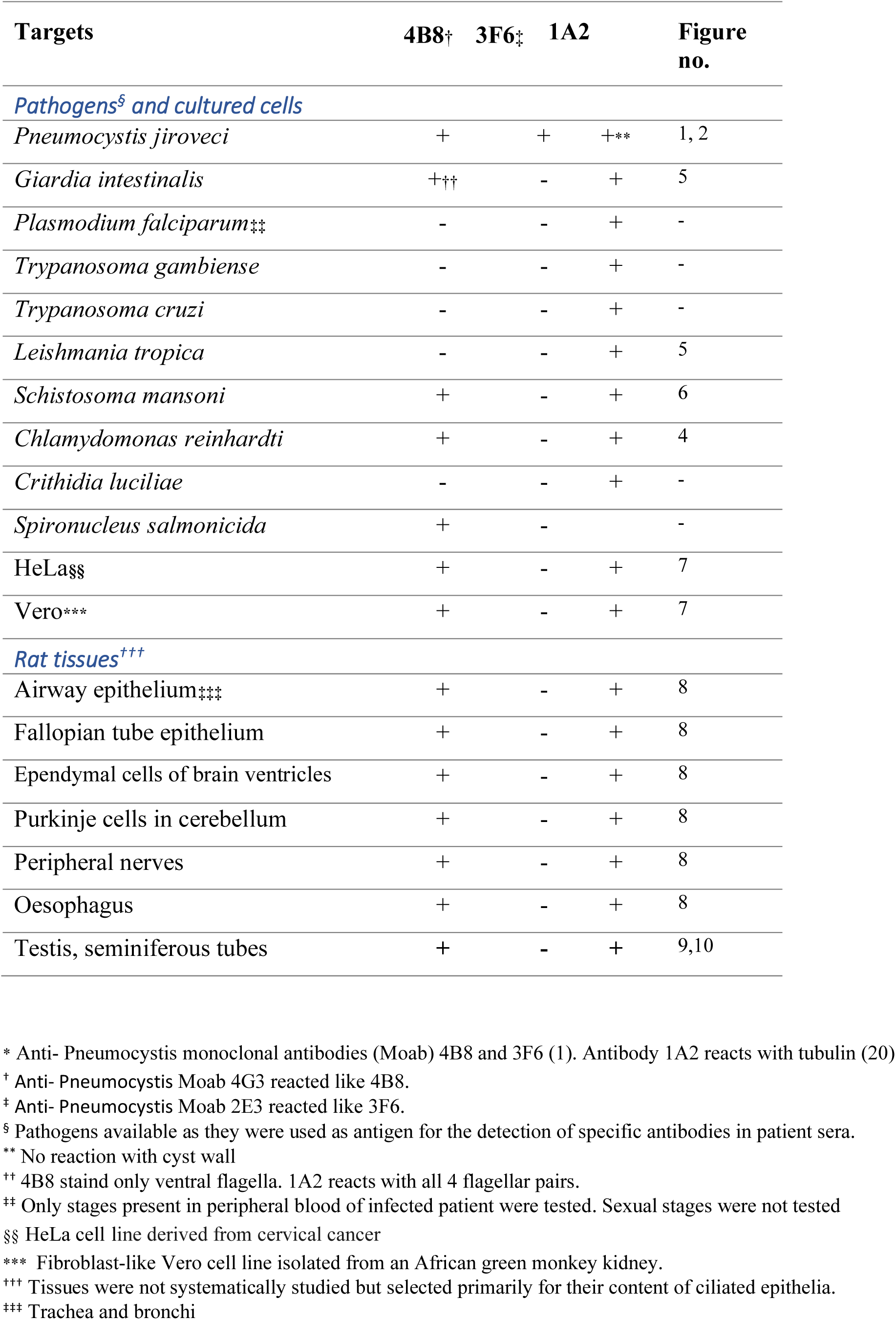
Reactivity of mouse monoclonal antibodies* with various targets in indirect immunofluorescence staining.

## Discussion

The surprising finding, that antibody 4B8 generated by immunization of mice with *Pneumocystis jiroveci* cross-reacts with cilia and is capable of distinguishing the ventral flagella of Giardia from the other flagella of this organism, provoked the question “what’s so special about the Giardia ventral flagella?” In trying to answer this question, an attempt is made to shed some light on host-pathogen interactions in Pneumocystis infection. Antibody 4B8, and the pneumocystis-specific antibody 3F6, reacted with intra-alveolar **clusters of Pneumocystis organisms** similarly. In such clusters, three main morphologically distinguishable structures are seen; asci, trophozoites and filopodia. The asci are easily recognized in light microscopy e.g after staining with silver stains (6)(22), but the vegetative stage, trophozoites, measuring a few μm in diameter and constituting 90-95% of the total parasite population in the lungs of infected hosts (23), are difficult to recognize in light microscopy without specific markers. (2)(24)(25)(26)(8)(21)(27) The third morphologically distinct morphological feature, tubular projections similar to filopodia or microvilli (28), can barely be recognized by conventional light microscopy (29), but surface projections (Fig 1) are seen as clusters hundreds of small tubules in ultrathin sections in both human and rat Pneumocystis in transmission electron microscopy. (30)(31)(32)(33)(34) The Pneumocystis-specific antibody 3F6 recognizes localized such tubular extensions in addition to both cysts and trophozoites. (35) Filopodia have been linked to nutrition of the organism (28)(29)(36), a function further suggested by the observation that filopodia are surrounded by tissue exudate, not by air. (29)(26)

Anti-tubulin antibody reacted with Pneumocystis trophozoites (Fig 2), which is consistent with the reported localization of tubulin in trophozoites at the ultrastructural level (37). Anti-tubulin antibodies stained clusters of organisms occupying the alveolar spaces in Pneumocystis pneumonia (Fig 2), but not the cyst wall, which is consistent with the ultrastructural observation by Bedrossian, who observed no microtubules in the cyst wall or tubular extensions. (38)

### Localization of the anti-4B8 target epitope in association with microtubules

Localization of the 4B8 target epitope was observed both at the tissue and the cellular level in association with cytoskeletal microtubules. This was evident from immunostaining both in cultured cells and in ciliated epithelia. The cytoskeletal microtubular network of cultured cells is easily recognized. Cilia/flagella, of e.g. Chlamydomonas algae have the common internal arrangement of 9+2 microtubule doublets, which constitute the conserved axonemal structure (39).

However, the 4B8 target epitope is not present in this basic axonemal structure as it is absent from some axonemes, such as flagella of Trypanosomes and interestingly lacking in 3 out of 4 flagellar pairs of Giardia. Interestingly all the axonemes if Giardia flagella appear to be similar: Reorganization of the flagellar apparatus in dividing Giardia involves **flagellar transformation**: “… a maturation process during which the flagella migrate, assume different position and transform to different flagellar types in progeny…” (40). This transformation through which the respective axonemes change their position takes place by reorientation and migration of the pairs of basal bodies. During the transformation process the flagellar function is altered as parent anterolateral flagella become caudal and both parent posterolateral and ventral flagella become anterolateral in daughter cells. The similarity of Giardia flagellar axonemes is consistent with the fact that no member of the tubulin family or post-translational tubulin modification is associated selectively with the Giardia ventral flagella. (see supplementary information in S1)

The anti-4B8 staining of the ventral Giardia flagella appeared to be located periphery of the axoneme proper and did not appear to stain the intracytoplasmic portion of the axoneme (see Fig 5). The pattern of staining of the Giardia ventral flagella was fine granular and appeared to be localized closer to the plasma membrane than the axoneme stained by anti-tubulin antibody 1A2.

Giardia species has been shown to contain numerous vesicular structures apparently forming a complex endomembrane system of protein sorting and transport which evolved early in eukaryotes (41)., Para-axial electron dense ancillary structures beneath the membrane that run alongside the axoneme from its point of emergence to the flagellar tip (corresponding to those seen in VF in Fig 11), and present only in association with the ventral flagellar axonemes have been demonstrated previously. (42)(43) These functionally uncharacterized paired paraflagellar structures characteristic of ventral flagella are dissolved after detergent treatment to produce pure axonemes for transmission electron microscopy. (44)(45)(46)(47)(48)

**Fig. 11.**
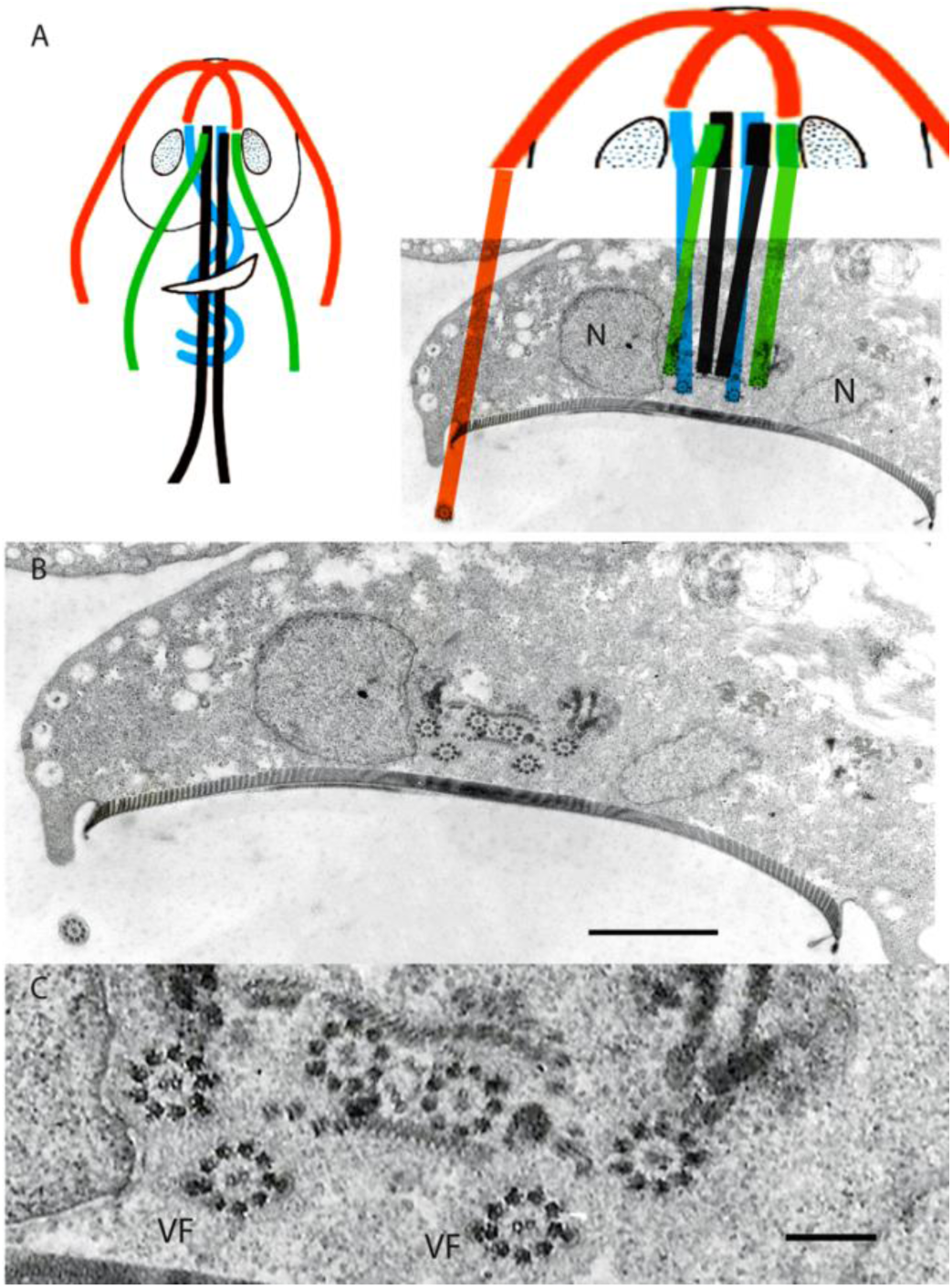
Transmission electron micrograph of Giardia trophozoite showing cross section of the four flagellar pairs running at the level of the caudal portion of the nuclei as indicated in the schematic drawing ***(A)*** (modified from (40)). Ventral flagella axonemes (blue and VF) are associated with two lateral electron dense masses. Antero-lateral flagella (red), postero-lateral flagella (green) and caudal flagella (black). Size bars 1 micron in ***B*** and 200 nm in ***C***.

Recent genomic and proteomic approaches have identified novel flagellar proteins and advanced our understanding of fundamental aspects of eukaryotic flagellum structure and function.(49). The number of components associated with the microtubules of the conserved 9+2 axonemal structure is surprisingly high, and their functional significance is only beginning to unfold.(39)(50)(51) In Giardia more than 300 microtubulus-associated proteins are thought to occur (52). The apparent localization of the 4B8 target in association with some axonemes is interesting as the axoneme has been shown to recruit distinct protein components to confer functional differences to cilia at distinct locations, e.g. mucus-propelling cilia in the trachea and water-propelling cilia in brain ventricles (53).

The subsequent discussion will consider to what extent the localization of the 4B8 target antigen is consistent with a conserved function such as intra-flagellar transport (IFT) (54).

### Localization in tissues

The 4B8 target epitope was seen in ependymal cells lining the brain ventricles and in Sertoli cells of the epididymis, two distinct tissues which share the functional characteristics of being part of the **blood-tissue barrier**. Ependymal cells lining of the brain are a type of glia cells which form a continuous sheet lining the ventricles, choroid plexuses and the central canal of the spinal cord. They have a simple columnar shape and are ciliated much like mucosal epithelia. Ependymal cells have a cerebrospinal fluid-brain barrier function and participate in the control of water transport and ion homeostasis.

In the seminiferous tubules of the testes microtubules are abundant in apical regions of Sertoli cells, and are closely related to ectoplasmic specializations attached to the adjacent spermatid heads. There is a close contact between Sertoli cells and mature germ cells (spermatids). Just prior to spermiation, the elongated spermatid interacts with the Sertoli cell via an extensive structure comprising various adhesion molecules called apical tubulobulbar complexes. (55)(56)(57)(58) Microtubule-based transport appears to occur both in the manchette and at intercellular adhesion junctions in the Sertoli cell plasma membrane. The cytoskeleton of Sertoli cells and the manchette of developing spermatocytes are known to contain abundant networks of microtubules. Structural proteins are apparently delivered to the basal body through intra-manchette transport and required for correct formation of the sperm tail and acrosome and shaping of the head. (59).

Cells of the central and peripheral **nervous system** react with antibody 4B8 both in the flatworm Schistosoma and in the rat. The parasitic schistosome worm belongs to the most primitive metazoan phylum, Platyhelminthes. The worm neuron is highly secretory and contains a heterogeneity of vesicular inclusions, dominated by vesicles, whose contents may be released synaptically or by paracrine secretion for presumed delivery to target cells via the extracellular matrix. (60)(61)

In the rat cerebellum, staining of large neurons located as a single layer between the outer molecular layer and the inner granular layer. These Purkinje cells are sole output neurons of the cerebellar cortex bridging motor and non-motor domains. Both peripheral and central nervous tissues are involved in vesicle transport and signal processing. **Axonal transport** of different cargos depends on an incompletely known set of molecules along intracellular tracks formed by the microtubule networks.(62)

The internalization of components of the plasma membrane-associated ligands and fluid is a fundamental process in eukaryotic cells and molecules involved in cliliary functions have apparently been present in a common ancestor about one billion years ago (63) **Membrane transport** is involved in diverse processes such as uptake of nutrients and intercellular signaling. Clathrin-mediated endocytosis is an essential cellular mechanism by which eukaryotic cells regulate their plasma membrane composition to control cell signaling, adhesion, migration and morphogenesis.(64) Attachment of Pneumocystis trophozoites to lung epithelium triggers proliferation and life cycle progression. (22) Interestingly, results of genome analysis show preservation in Pneumocystis of nearly all proteins associated with clathrin-dependent endocytosis. (32) Wether membrane transport occurs through the vast surface area provided by Pneumocystis tubular extensions or filopodia needs to be shown, but a plausible working hypothesis is that Pneumocystis filopodia are involved in **uptake of nutrients** provided by the host. Such a function is consistent with the observed location of the anti-4B8 target antigen in Chlamydomonas flagella and the Giardia ventral flagella, which are known to be involved in flagellar functions unrelated to the flagellar beat. The region between the flagellar plasma membrane and the outer tubulins of the axoneme is a highly complex structure where trains of particles located closely to the inner surface of the flagellar membrane bridge to each other and to the outer doublet microtubules.

(65)(42)(66)(67) Intraflagellar transport (IFT) of cargoes takes place along the length of the flagella along microtubules and the components of IFT and their assembling in the flagella seem to be highly conserved during evolution and their encoding genes also exist in the Giardia genome. (65)(68)(69)(70)

A closer look into the protein components of the Giardia cytoskeleton shows that it contains mainly **giardins** in addition to tubulins. (71) The multi-gene giardin family has 21 members (72), which are associated with Giardia flagella differently (73); The *α*-13 giardin is a cytoplasmic protein (74), the α18-giardin localized at all flagella (74), and the α-11 giardin is mainly localized to the plasma membranes and basal bodies of the anterior flagella (75). A striking observation is that **α-19-giardin is found only in the Giardia ventral flagella** (76): Polyclonal antibodies raised against a recombinant α-19-giardin protein reacted in a Western blot assay with a protein of about 47 kDa in the pellet fraction of a Giardia trophozoite extract. (76) Thus, both the localization of the 4B8 target epitope to the ventral flagella and its molecular mass of about 50kDa raises the possibility, that antibody 4B8 recognizes α-19-giardin.

The α-giardins are recognized as **annexin** homologues based on sequence similarities and in the novel nomenclature, Giardia annexins have been given the designation, annexin E. Annexins are Ca2+ and membrane binding proteins forming an evolutionary conserved multigene family comprising >500 different gene products expressed throughout animal and plant kingdoms. Many of the group E annexins interact with different cytoskeletal structures or the membrane. Annexins are involved in numerous cell processes including vesicle trafficking, calcium signaling, cell growth, division, and apoptosis (77)(78). Annexin A13b is known to interact with other proteins and helps to direct them to their destination points. In epithelial cells, this annexin is suggested to play a role in the formation of transport vesicles and their fusion with the apical membrane (79). It is conceivable that annexins exist in Pneumocystis as they are found in most medically important fungal pathogens. (80) (81) A final point for discussion is the evidence for an immune response selectively against Giardia ventral flagellar antigens and their possible **involvement in infection immunity**. Interestingly detergent soluble molecules selectively localizing to the ventral flagella appear to be immunogenic and to induce a humoral immune response in both human and murine Giardia infections. (43) Wether protective immunity against Giardia infection mediated by mothers milk (82) involves antibodies against the Giardia ventral flagella - and the 4B8 target antigen - is not known. However, preliminary results (83)(84) showed that 42% (25 out of 59) of mothers milk samples from a giardiasis-endemic region, contain antibodies reacting selectively with Giardia ventral flagella. This was twice the prevalence of antibodies reacting with all Giardia flagella in this material.

Thus, the results of this study suggest that looking for more answers to the question “What’s so special about Giardia ventral flagella?” may reveal novel aspects of host-pathogen interactions.

## Acknowledgements

Several persons have kindly contributed over the years to the experiments resulting in this report and the results could not have been obtained without their generous help. To name the most important contributors, Cecilia Thors† performed much of the laboratory work together with Jadwiga Winiecka-Krusnell who managed to rescue monoclonal antibodies through the turmoil of institutional re-organizations. Protozoa and helminth materials were obtained from the diagnostics laboratory of the Parasitology Department of the Swedish National Bacterilology Laboratory (SBL). Carl-Henrik von Bonsdorff performed the electron microscopy. Thomas Kreis† provided anti-tubulin antibodies and Margareta Wallien the bovine brain tubulin preparation. The Chlamydomonas culture was a gift from Susan Dutcher and the Spironucleus specimens were from Staffan Svärd.

Johan and Mikael Lundin arranged for supplementary image material to be accessible at the Webmicroscope.net website. My former students Seppo Meri and Antti Sukura are thanked for encouragement.

## S1 Discussion on Giardia tubulins

As essential components of the eukaryotic cytoskeleton, microtubules fulfill a variety of functions that can be temporally and spatially controlled by tubulin posttranslational modifications. (63)(85)(86) The observation that the 4B8 target antigen has a molecular size similar to beta tubulin and a localization in the microtubular cytoskeleton suggested that the it may be a novel tubulin isoform with a mobility similar to beta tubulin (87). However, no tubulin isoform or posttranslational modification associated only with ventral flagella is known. The axonemes of ventral flagella are involved in “… a maturation process during which the flagella migrate, assume different position and transform to different flagellar types in progeny…” (40). This transformation is mediated by reorientation and migration of the pairs of basal bodies through which the respective flagella change their position and function. During the transformation process the parent anterolateral flagella become caudal, whereas both parent posterolateral and ventral flagella become anterolateral in daughter cells. This similarity of Giardia flagella is consistent with the fact that no member of the tubulin family or post-translational tubulin modification is associated selectively with the Giardia ventral flagella Despite the immunolocalization of antibody 4B8 to microtubules in cultured cells and cilia/flagella and reaction with a component of about 50kDa in western blotting, the 4B8 target epitope does not seem to be any known tubulin. Its presence in semi-purified preparation of bovine brain tubulin suggested initially that the antigen epitope was related to beta tubulin. However, by preparing purified tubulin from bovine brain by cycles of assembly and disassembly, the preparation contains microtubule-associated proteins(88). The target could be some specific epitope related to post-translational tubulin modification or a member of the tubulin family other than the tubulins assembled into 9+2 protofilaments of the microtubule wall (89). However, no reported tubulin (classes γ, δ, ε, η, ζ and ι) exhibit a selective reactivity with Giardia ventral flagella like the one observed with antibody 4B8 (44)(45)(90)(91)(92). Also no selective reactivity with ventral flagella was reported in studies on a panel of 23 monoclonal antibodies against different epitopes on N and C termini of alpha- and beta-tubulins and against known posttranslational modifications of tubulin.(40) This was confirmed in our experiments using a number of antibodies against tubulin-related antigens(93): These antibodies (from Sigma, St. Louis, USA) include mouse monoclonal antibodies reacting with alpha tubulin (clone B-5-1-2); anti-beta tubulin (clone tub 2.1); anti-delta tubulin (antibody T3950 clone DTU-64) anti-epsilon tubulin (antibody T1323 clone TUB-11); anti-gamma tubulin (antibody T6557 clone GTU-88); anti-acetylated tubulin antibody (T6793 clone 6-11B-1); anti-polyglutamylated tubulin antibody (T9822 clone B3) and the polyclonal antibody to bovine brain microtubulus-associated proteins recognizing MAP2 and to a lesser extent tau (94).

